# Alteration of the social and spatial organization of the vector of Chagas disease, *Triatoma infestans*, by the parasite *Trypanosoma cruzi*

**DOI:** 10.1101/383620

**Authors:** Stéphanie Depickère, Gonzalo Marcelo Ramírez-Ávila, Jean-Louis Deneubourg

**Author notes:** Corresponding author: Stéphanie Depickère.

## Abstract

Insects of Triatominae subfamily are vectors of the parasite *Trypanosoma cruzi*, the etiological agent of Chagas disease affecting millions of people in Latin America. Some of these vector species, like *Triatoma infestans*, live in the human neighborhood, aggregating in walls or roof cracks during the day and going out to feed on animal or human blood at night. Except for their feeding specialization, these insects share this cycle of activities with many gregarious arthropod species. The understanding of how sex and *T. cruzi* infection affect their aggregation and geotaxis behavior is essential for understanding the spatial organization of the insects and the parasite dispersion. Experiments with non-infected and infected adults of *T. infestans* show that the insects presented a high negative geotaxis and aggregative behavior. Males had a higher negative geotaxis and a higher aggregation level than females. The aggregation level and the negative geotaxis were stronger in infected insects than in non-infected ones, the difference between sexes being maintained. The importance of these results is discussed in term of parasitic manipulation, dispersion of the vector and strategy of its monitoring.

## Introduction

Chagas disease is one of the most important neglected tropical diseases with 6-7 million people who are estimated to be infected, and 20% of the population who are at risk^1–3^. This vector-borne disease is caused by the parasite *Trypanosoma cruzi* (Kinetoplastida: Trypanosomatidae) and is mainly transmitted by contact with infected feces/ urine of hematophagous insects of the Triatominae subfamily (Hemiptera: Reduviidae). Currently, 149 extant species have been described worldwide, and all of them are considered as potential vectors^4^. Most of them live in sylvatic habitats, and only a dozen of species are regarded as vectors of major epidemiological importance due to their capacity to live in the surrounding of the human dwellings where they find stable shelters and food abundance^5^. *Triatoma infestans* is the main vector in the Southern Cone of South America. Except for their feeding specialization, the domiciliary species share similar lifestyle and cycle of activities with many gregarious arthropods including other synanthropic species like cockroaches^6–8^. During the daytime, they assembled in dark and sheltered places such as cracks in the walls or roof, or behind objects hanging on walls. At night, they leave their shelter to actively seek a host upon which to feed and then, they come back to a resting place to digest. The digestive phase can last from some days to several weeks according to the blood meal size, the individual and the environmental conditions^9^.

The control strategy for Chagas disease relies mainly on the control of the domestic vectors through chemical control^1^. Faced with the increased of the insecticide resistance exhibited by these insects, and with the reinvasion of the dwellings by residual or sylvatic population of triatomines^10–12^, it is necessary to study the behaviors leading to a better understanding of their distribution and their dispersion. In this perspective, aggregation and geotaxis are key behaviors. Knowing them better and understanding how the parasite dispersion may influence them is fundamental. Indeed, aggregation is a widespread behavior that results from a response of individuals to environmental heterogeneity, and from social interactions involving attractions between individuals^13–15^. The social interactions maintain the group cohesion and the associated adaptive values of group living. In triatomines, protection against predation is usually evoked as the main benefice of clustering, but surviving might also be enhanced thanks to protection against hydric loss, and to a higher probability of coprophagy, symbiont exchange, and of sex encounters, as it was shown for other insects^16–20^. Aggregation in triatomines was investigated with a focus towards the substances that mediate it, and on the factors that modulate the aggregative response^21–26^. All these works analyzed nymphal instars behavior response; in adults very few is known except that they can aggregate around feces^25^. Geotaxis, also called gravitaxis, is a crucial behavior involved in insect orientation^27^. Animals can exhibit locomotion that is gravitationally directed vertically down or up (positive or negative geotaxis, respectively). Geotaxis in triatomine has been poorly described, *T. infestans* was just reported as being more concentrated in the upper half of the walls in houses or chicken houses^17,28^. Moreover, to our knowledge, no studies were conducted to analyze the synergy or conflict between gregariousness and geotaxis in triatomines.

It is well-known that parasites can modify physiological, behavioral, and/or morphological traits of their hosts to increase their fitness, even if it is at the cost of the host fitness^29^. The latter usually means that infected hosts will behave in ways that facilitate the transmission of the parasite^30,31^. Literature about the effects and possible manipulation of triatomines behavior by *T. cruzi* is relatively sparse, covering only seven species: *Mepraia spinolai, Panstrongylus megistus, Rhodnius pallescens, Rhodnius prolixus, Triatoma brasiliensis, Triatoma dimidiata* and *T. infestans*. Authors have been especially interested in the parasite’s effects on four groups of the host’s behavior: life-history trait, feeding, defecation, and dispersion/ locomotion. It seems that *T. cruzi* increases the development time and biting rate, and decreases the longevity and defecation time in *M. spinolai*^32,33^, but no change was observed in *P. megistus*^34^, *R. prolixus*^35^, *T. dimidiata*^36^, *T. infestans*^37^, and almost no change in *T. brasiliensis*^38^. The reproduction was decreased by *T. cruzi* in *T. brasiliensis*^38^. The dispersion was higher in infected females of *T. dimidiata* than in non-infected females; no effect was found in males^39^. Moreover, *T. dimidiata* individuals infected with *T. cruzi* were found to have larger wings than non-infected ones^40^. In *R. pallescens, T. cruzi* infection did not significantly impact flight initiation, but it was observed that infected females flew significantly faster than males from 30 s to 2 min after flight initiation^41^. The locomotory activity of *R. prolixus* was decreased by infection: the total number of movements was 20% less than that observed in non-infected insects^42^. The time to find a host for an infected *M. spinolai* was almost twice as fast as for a non-infected insect^33^. In conclusion, modification of the triatomine traits seems to be species-dependent, age-dependent, sex-dependent, and even environment/ physiology-dependent.

In this work, video-recorded experiments were conducted to study aggregation and geotaxis in adults of *T. infestans* and to analyze the effect of the infection with *T. cruzi*. Our hypotheses, based on the literature, were that these two behaviors – gregariousness and geotaxis - are strongly intertwined and are increased in infected individuals. In each experiment, ten insects (non-infected females, non-infected males, infected females, or infected males) were dropped at the base of a vertical wall covered with a paper sheet allowing the bugs to climb. Spatial positions of each insect were extracted from the video every five minutes until 150 minutes, permitting the following of the dynamics and the calculation of the size and spatial stability of the clusters. We demonstrate that both sexes exhibit a high clustering and a high negative geotaxis, males revealing a higher response than females. Interestingly, the *T. cruzi* infection significantly strengthens both behaviors in both sexes.

## Results

### Negative geotaxis: spatial distribution and total population

The bugs quickly climbed on the wall and stayed there, demonstrating a high negative geotaxis; and after an exploratory phase, the insects began to cluster and rest (see Supplementary Fig. S1 online). After 10 min, more than 80% of the individuals were on the wall (90% after 20 min) for the four conditions; and this proportion remained constant until the end of the experiment where no statistical difference was detected between the four conditions (Fig. 1). The bugs were mostly located in the upper half of the setup; the median vertical position reached a plateau value (stationary state) after 15 min with a value greater than 35 cm for the four conditions (Fig. 2). Their vertical distributions at 150 min (end of the experiment) revealed a statistical difference between sexes, e.g., males were located higher than females, and also between non-infected and infected individuals, e.g., infected males were higher than non-infected males (Fig. 3). These trends were also discovered inside the top strip of 4 cm (40-44 cm), a zone corresponding to 10% of the total area of the setup and where 56% (80%) of the non-infected (infected) males and 33% (44%) of the non-infected (infected) females were located (see Supplementary Fig. S2 online).

**Figure 1.**
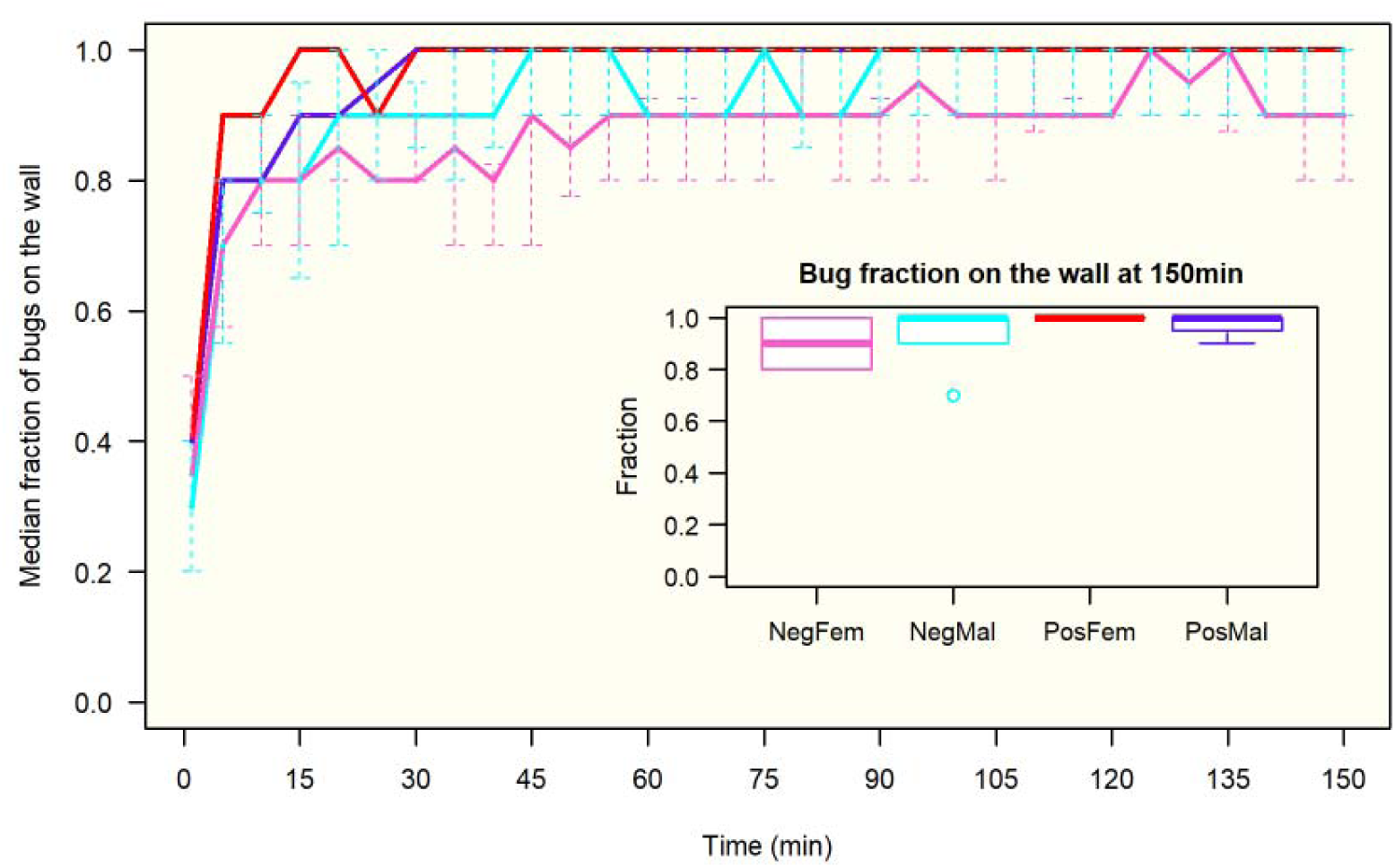
Median fraction of bugs on the wall (quantiles 25%-75%). Number of experiments: 16, 9, 12 for NegFem (pink), NegMal (cyan), PosFem (red) and PosMal (blue) respectively. Inserted figure: boxplot distribution of bugs on the wall at 150 min. Anderson-Darling test at 150 min between the 4 conditions: TkN = −0.12, P = 0.46.

**Figure 2.**
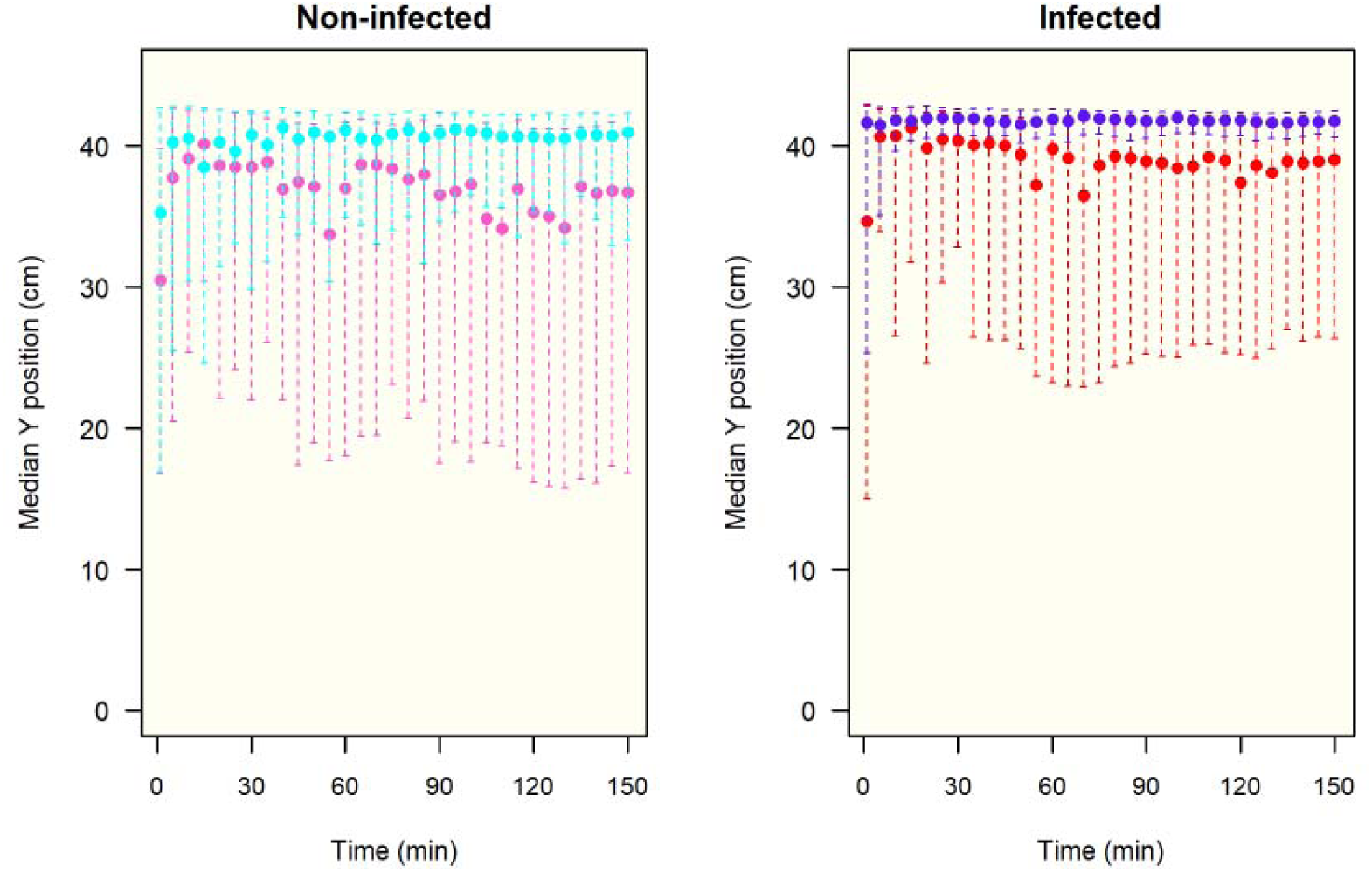
Evolution of the median vertical position (quantiles 25%-75%) of the non-infected and infected insects on the wall during the experiment. Median position at 150 min: NegFem (pink): 36.7 (16.8-41.2) cm, PosFem (red): 39.0 (26.3-42.0) cm, NegMal (cyan): 40.9 (33.3-42.3) cm, PosMal (blue): 41.8 (40.6-42.5) cm.

**Figure 3.**
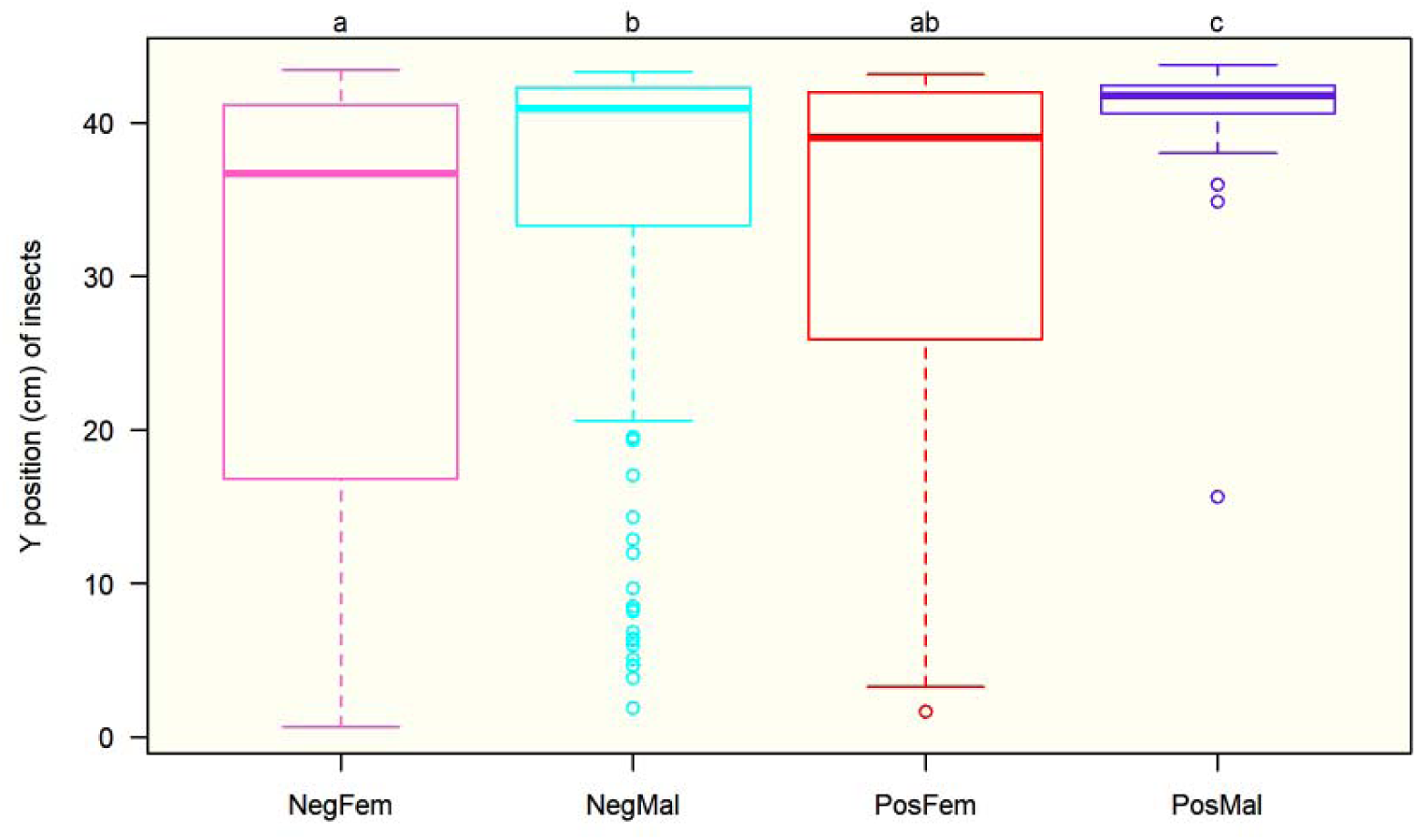
Boxplot distribution of the vertical position of the individuals at 150 min. Number of observations: non-infected females: 145; non-infected males: 141; infected females: 90; infected males: 117. Anderson-Darling k-sample test between the four conditions for all individuals: TkN = 24.4, P < 0.001. Results of Anderson-Darling all-pairs comparison tests are shown at the top of the figure (conditions with different letters correspond to conditions statistically different at P < 0.001).

At the end of the experiment, more than 80% of the individuals had a vertical orientation from which around 70-80% had the head turned towards the top of the setup (± 30°). For the four conditions, individuals were not uniformly distributed (Rao’s test < 0.01), but rather centered on 0 (V-tests < 0.001, Fig. 4). When the distributions of individual orientations were compared, no difference appeared between sexes. Interestingly, the infection affected the orientation of the males which demonstrated a higher proportion of insects with the head towards the bottom when infected (Fig. 4).

**Figure 4.**
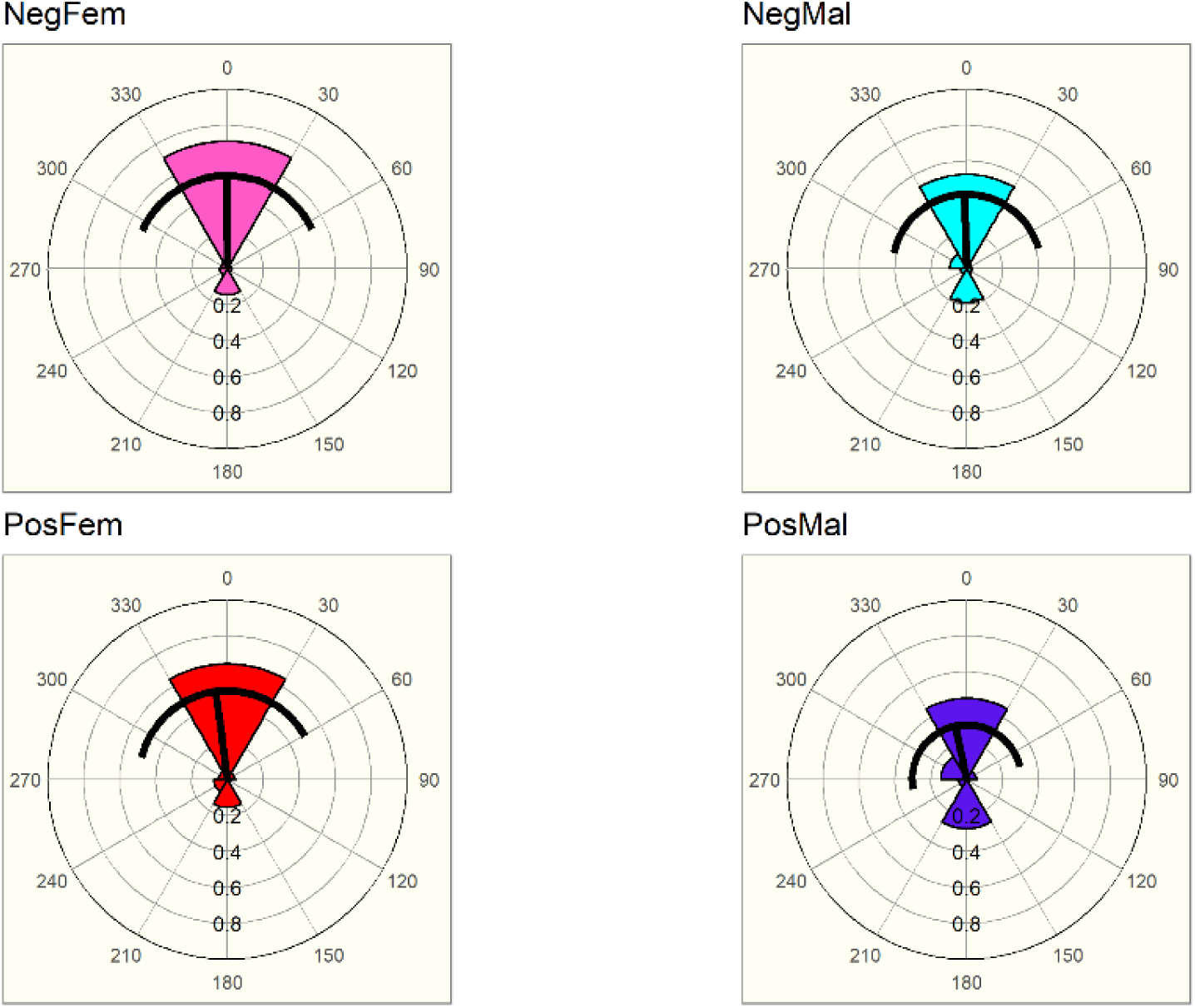
Vertical orientation of insects for the four conditions. Wedge’s angles of 60°, frequencies are shown as radius of the wedge. 0° represents the head to the top. The dark line shows the mean direction and its length, and the standard deviation of the distribution. Number of observations: NegFem: 145, NegMal: 141, PosFem: 90 and PosMal: 117. Rao’s tests gave P < 0.01 for the four conditions. V-tests (testing the null hypothesis of uniformity against non-uniform distribution with a mean of 0) gave P < 0.001 for the four conditions. Mardia-Watson-Wheeler pairwise tests between the four conditions: NegFem/ NegMal: P = 0.24, NegFem/ PosFem: P = 0.156, NegFem/ PosMal: P = 0.001, NegMal/ PosFem: P = 0.033, NegMal/ PosMal: P = 0.005, PosFem/ PosMal: P = 0.049.

To summarize, without being infected, both sexes exhibited a high negative geotaxis that was higher for males than for females (Fig. 3). Most of the insects of both sexes were oriented the head toward the top (Fig. 4). The *T. cruzi* infection strengthened this bug’s geotaxis, especially for males.

### Global Clustering

The median aggregated bug fraction increased up to reach a plateau around 35 min, gathering around 70% and 90% of non-infected and infected males respectively, and 40% and 60% of non-infected and infected females respectively (Fig. 5). A statistical difference was detected at 150 min between sexes (males showed a higher aggregated fraction than females), and between infected conditions (infected bugs with a higher aggregated level than non-infected ones) (Fig. 5, see also Supplementary Fig. S3 online). Insects in all conditions tended to gather in one or two clusters. The biggest cluster assembled 40% (70%) of the aggregated non-infected (infected) population in males, and 30% (40%) of the aggregated non-infected (infected) population in females (Fig. 5). No difference was observed between sexes, neither between infection condition (Fig. 5). When the structure of the clusters was compared between infected sexes, clusters of infected males looked more compact, with a significantly smaller distance between aggregated individuals; and they also looked denser, with a higher K-density (Fig. 6, see also Supplementary Fig. S4 online).

**Figure 5.**
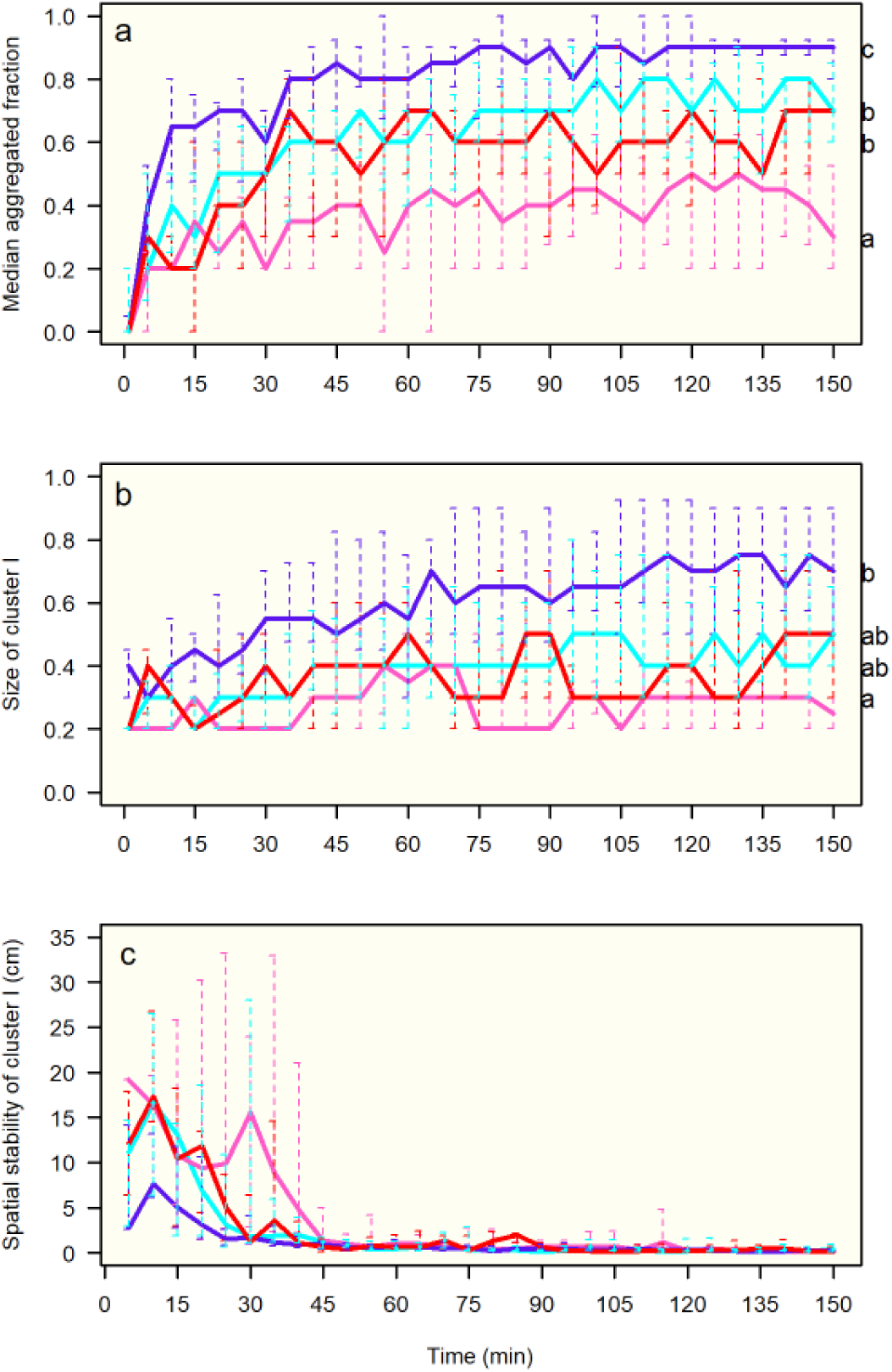
Dynamics of aggregation for the four conditions: NegFem (pink), NegMal (cyan), PosFem (red) and PosMal (blue). (a) median fraction of aggregated individuals (quantiles 25% - 75%); (b) median size of the biggest cluster (quantiles 25% - 75%); (c) spatial stability of the biggest cluster (quantiles 25% - 75%). Anderson-Darling k-sample test at 150 min between the four conditions: (a) TkN = 7.4, P < 0.001, (b) TkN = 4.9, P = 0.001, and (c) TkN = −0.5, P = 0.63. Results of Anderson-Darling all-pairs comparison test are shown with different letters corresponding to conditions statistically different at P < 0.05 (on the right side of the figure).

**Figure 6.**
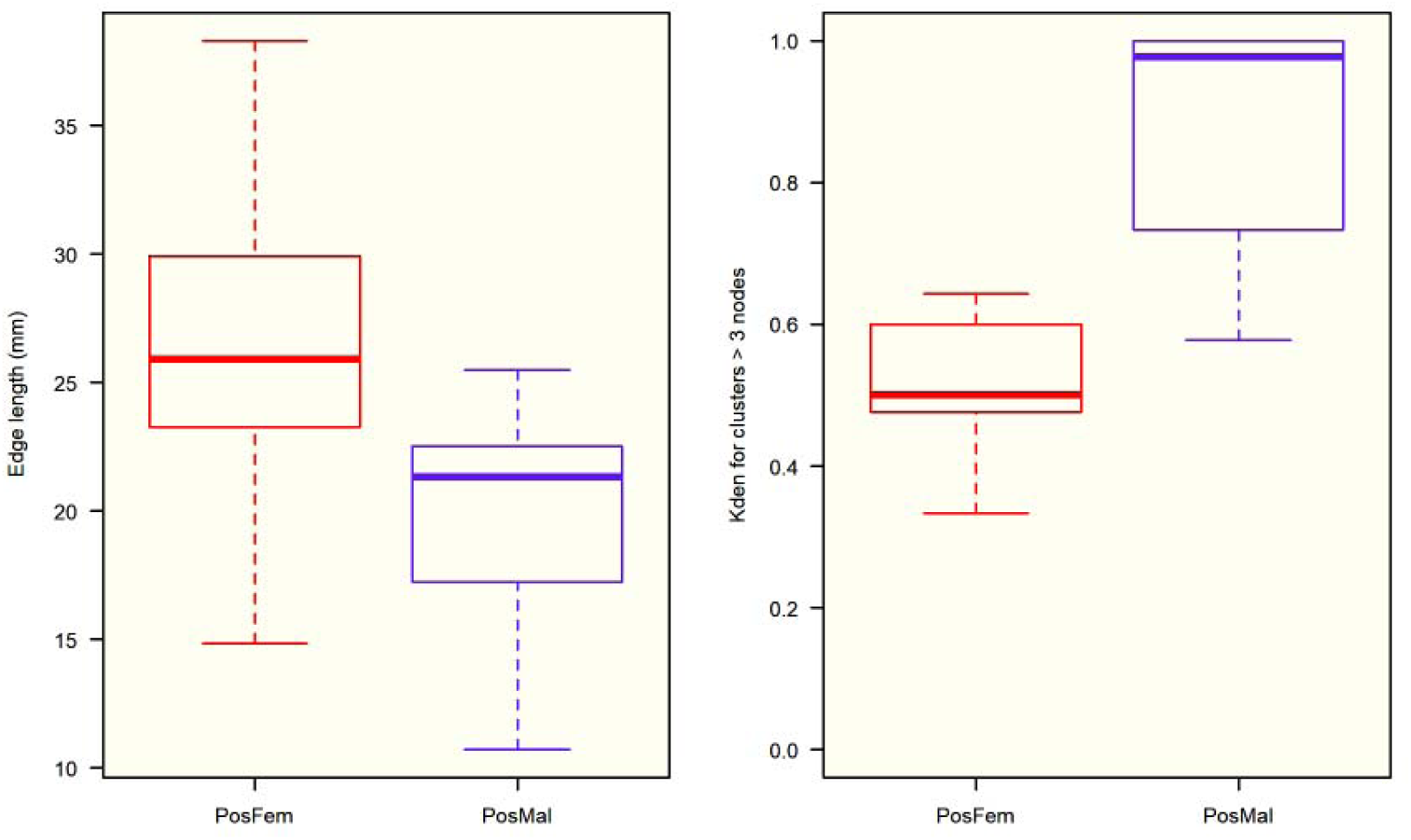
Structure of the clusters in infected conditions. Boxplot of the size of edges (or links) between aggregated bugs (left) and K-density of the clusters of size > 3 individuals (right). Anderson-Darling k-sample test for size of the edges: TkN = 9.1, P < 0.001 (72 and 255 observations for PosFem and PosMal respectively); and for the K-density: TkN = 4.7, P = 0.004 (17 and 21 observations for PosFem and PosMal respectively).

At the end of the experiments, individuals were very stable in space: the median fraction of individuals that moved less than 1 cm was greater than 60% for the four conditions, and no difference between sexes and infection condition was detected (Fig. 7). The biggest cluster also showed a strong spatial stability, with no statistical difference between the four conditions (Fig. 5).

**Figure 7.**
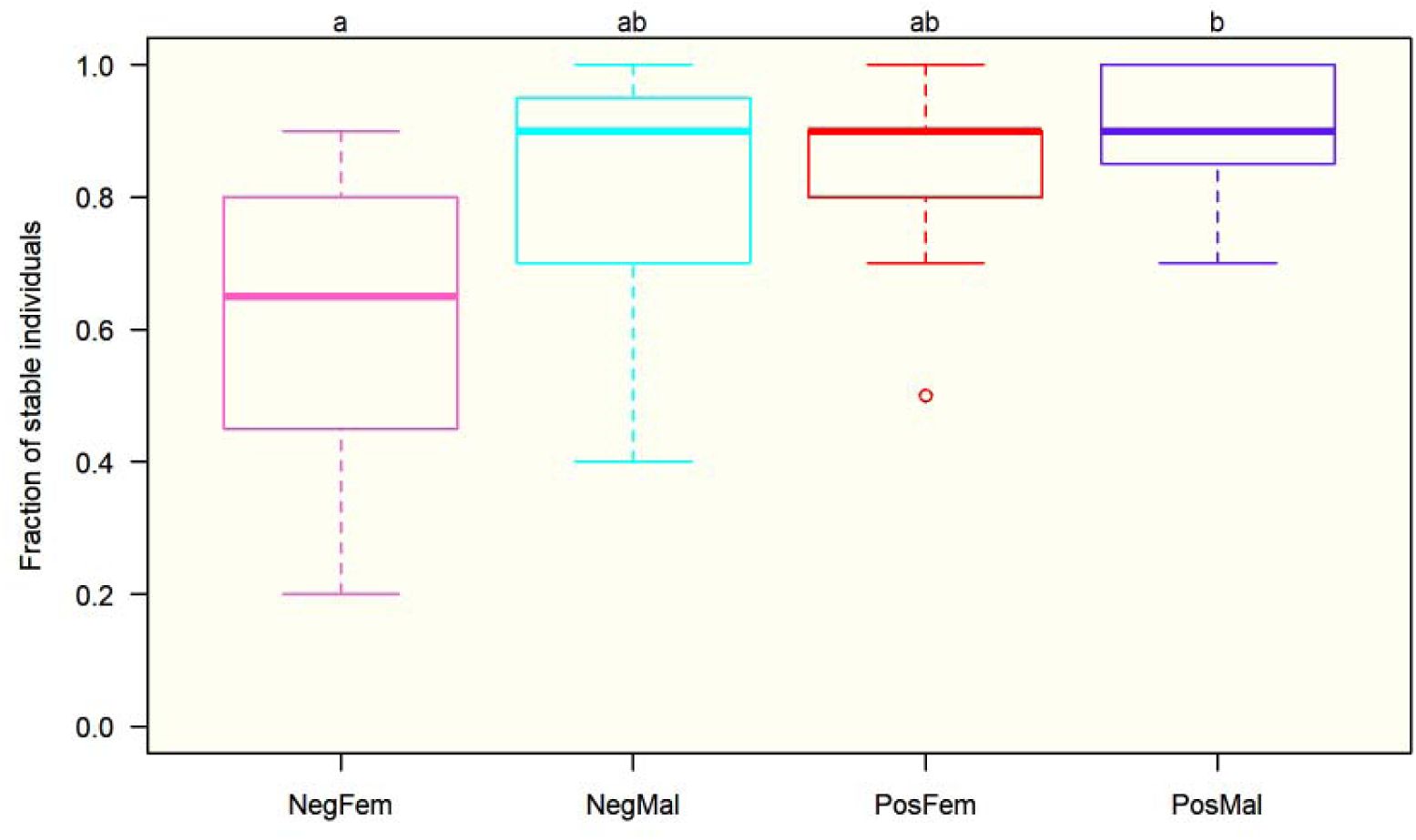
Boxplot of stable individuals (move < 10 mm between two snapshots) for the four conditions. Anderson-Darling k-sample test between the four conditions: TkN = 2.8, P = 0.017. Results of Anderson-Darling all-pairs comparison tests are shown at the top of the figure (different letters correspond to conditions statistically significantly different at P < 0.005).

In order to verify that the fraction of aggregated individuals was not directly due to the method of calculating this fraction and the increase of the bugs density at the top of the setup, 20,000 repetitions of groups of *N* simulations were performed (N = 16 for NegFem, N = 15 for NegMal, N = 9 for PosFem, and N = 12 for MalPos). For each simulation, 10 points were vertically distributed following the experimental vertical distribution of the bugs at 150 min (see Supplementary Figure S5 online), and homogeneously horizontally distributed. For each group of simulations, the mean fraction of aggregated individuals was calculated for each repetition. The mean aggregated fractions obtained in the simulations were 0.26, 0.42, 0.36, and 0.72 for NegFem, NegMal, PosFem, and PosMal respectively, revealing that an increase of the geotaxis leads to a rise in the observed aggregation level. However, the probability of observing a mean aggregated fraction higher or equal to the corresponding experimental one was P < 0.0001 for all the conditions, demonstrating that the observed phenomenon involved an active aggregation due to the inter-attraction between individuals.

The fraction of aggregated individuals in a strip of 0.5cm was proportional to the fraction of the population settled in this strip (Fig. 8). The slope of the regression line was the lowest for the non-infected female condition, and the highest for the infected male condition, being intermediate and similar for the two other conditions. The slopes of the linear regression were compared computing a model including the interaction between the total number of bugs and the conditions: a significant interaction was found (F_3, 348_ = 30.04, P < 0.001), giving a slope equal to 0.61 for non-infected females, 0.85 for non-infected males, 0.84 for infected females, and 0.92 for infected males. The comparison of the slopes between the four conditions showed that the non-infected females’ slope was lower than the three other conditions’ slopes (P < 0.001), the infected males’ slope was higher than the three other conditions’ slopes (P < 0.02), and the non-infected males’ slope was not different from the infected females’ slope (P = 0.98). These results demonstrated that, for the same density, the aggregation was higher for males than for females and for infected insects than for non-infected ones.

**Figure 8.**
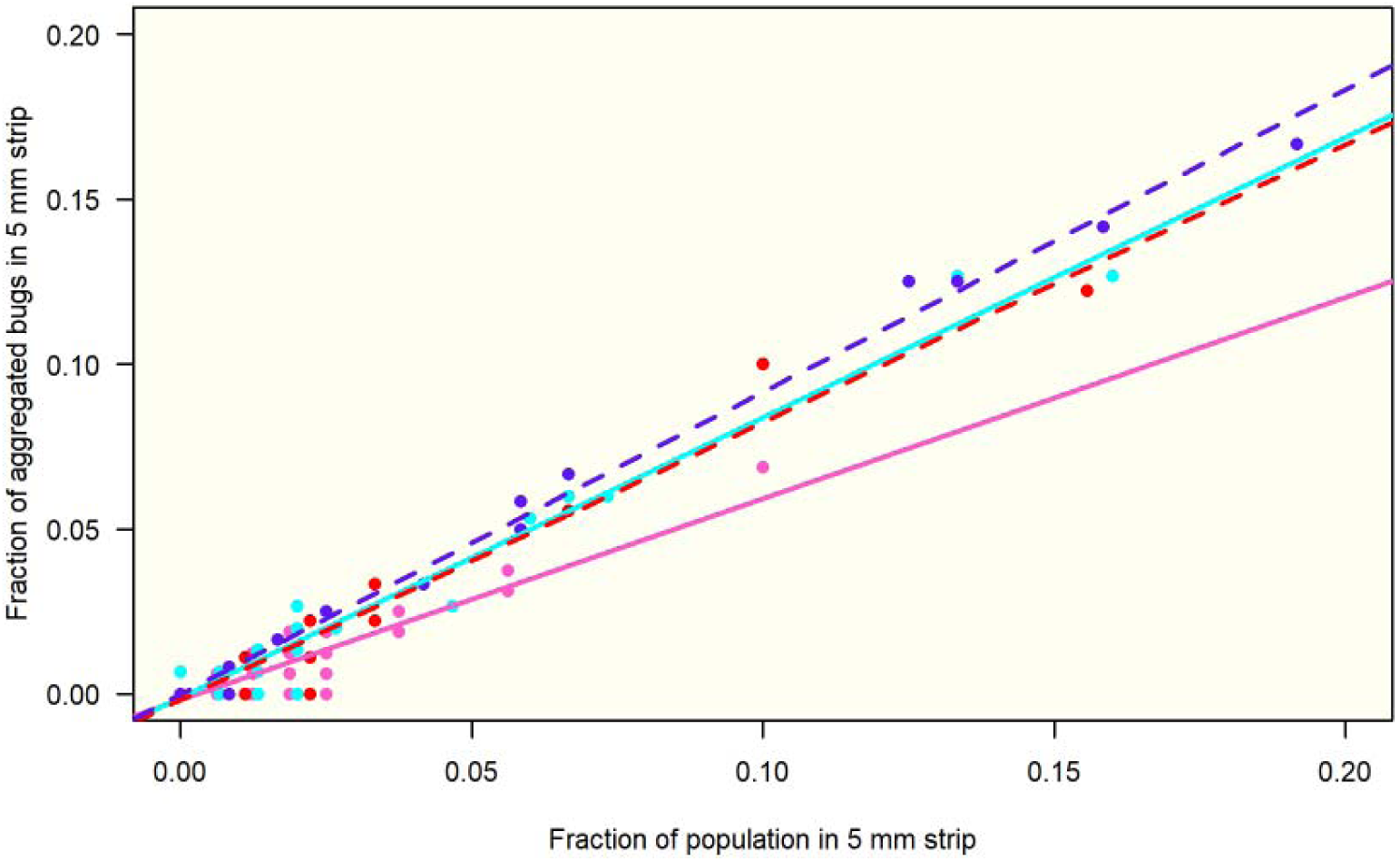
Fraction of aggregated individuals according to the population present in a horizontal strip of 5 mm high for the four conditions. Linear regressions: NegFem (pink): y = 0.6106 × - 0.0019 (SE of slope: 0.0296), R^2^ = 0.83, P < 0.001; NegMal (cyan): y = 0.8498 × - 0.0013 (SE of slope: 0.0180), R^2^ = 0.96, P < 0.001; PosFem (red): y = 0.8408 × - 0.0017 (SE of slope: 0.0269), R^2^ = 92, P < 0.001; PosMal (blue): y = 0.9168 × - 0.0001 (SE of slope: 0.0076), R^2^ = 0.99, P < 0.001.

## Discussion

This work represents the first detailed analysis of aggregation and geotaxis in adult males and females of *T. infestans*, and how both sexes are affected by *T. cruzi* infection. As shown before in nymphal instars^43,44^, adults exhibited an active aggregation due to the inter-attraction between individuals, illustrating the social character of these insects. A stable aggregation emerged for both sexes, but the fraction of aggregated individuals and the density of the clusters were higher for males than for females. This difference between genders was maintained under *T. cruzi* infection, but the latter reinforced the gregariousness in both sexes.

Our results are in agreement with those of previous studies. Indeed, a multi-factorial analysis (using species, development stages and, feces source altogether) of the aggregative response of individuals to feces shows that the aggregation level was lower (but not statistically different) for females than for males^25^. It is well established that clustering or reduction of the inter-individual distances of social and subsocial/ presocial arthropods reduces various stresses and therefore energy consumption^18,45,46^. Different studies show that clustering reduces water loss^18–20,46^. We hypothesize that the clustering of *T. infestans* individuals provides a similar benefit. As their weight is lower than females, males could be under higher hydric stress, leading them to a stronger aggregation. Moreover, it could be more adaptive for females to aggregate less to disperse their eggs and increase their probability of survival. In our experiments, males and females were supposed to be in similar physiological status due to their comparable period of starvation (8-10 days), but in the case of infection, *T. cruzi* and *T. infestans* compete for nutrients, and bug individuals show reduced resistance to starvation when they are infected^47^. It might be speculated that infected bugs were more starved and therefore exhibited a stronger aggregation to reduce the cost of the different stresses.

We know very little about the distribution of triatomines inside a dwelling. Even if all the stages are found in the upper part of the walls^17,28^, our results suggest that males and females rest in a stratified manner on the walls, males being above of females. Domestic cockroaches have a similar way of life than triatomines, except the feeding habits. *Periplaneta americana*, for instance, shows a preference for vertical areas, and males were vertically positioned above females^48^. Due to the vertical air current, they can detect the female pheromone easily and orient themselves towards them. In triatomines, the sex pheromone is emitted by the female metasternal glands, inducing males moving towards the females (positive anemotaxis)^9^. On a wall, due to the difference in temperature between the bottom and the top (up to 6°C ^17^), an air current move towards the top of the wall could allow the males to feel at some distance the sexual pheromone released by the females. Most individuals in this study showed a vertical orientation with the head towards the top, allowing them to escape quickly from predators generally located below when positioned vertically on a wall. *T. cruzi* infection enhanced negative geotaxis, especially in males, where a higher proportion of infected individuals faced their heads toward the bottom of the experimental wall.

Detection of *T. cruzi* by direct microscopic observation is known as being less efficient than by polymerase chain reaction (PCR), especially in case of low parasitemia^49,50^. It is consequently possible that a small proportion of non-infected insects would have been detected as infected by PCR. Despite this handicap, the difference observed between groups was statistically significant. In the same way, a part of the experiments with non-infected insects was composed of a mix of non-infected and infected insects. The latter implies that a small proportion of infected individuals inside a group of non-infected bugs is not enough to observe a change at the group level. Another interesting question is whether all the discrete typing units (DTUs) of *T. cruzi* and even strains of these DTUs, will influence the behavior of the bugs in a similar manner. This point was recently asked by Peterson et al., demonstrating that different strains of *T. cruzi* had different consequences in life history outcomes of *R. prolixus*^51^. Thus, more experiments are necessary to understand how *T. cruzi* affects the mechanisms underlying the geotaxis and the clustering, from a physiological and a behavioral point of view.

Behavioral alterations upon infection are called parasitic manipulation when they are adaptive for the parasite, altering phenotypic traits of its host in a way that enhances its probability of transmission. Some examples where the parasitism affects the geotaxis and the gregarious behavior of the hosts were described^52-57^. Is there any advantage to *T. cruzi* to enhance the negative geotaxis and the aggregation behavior in males of *T. infestans*? Two hypotheses can be put forward: 1) expand its spread, by increasing the longevity of the vector, away from ground predators, and/ or by allowing a higher rate of coprophagy and cleptohaematophagy through a stronger aggregation; and 2) facilitate the host-finding; it could be easier for them to find a food source by being higher on a vertical surface. Actually, these insects are known to fall near a host from the ceilings when they wander in a host search behavior^58^. Ramirez-Sierra et al. (2010) have already reported an increase of the dispersion on the field of infected females of *T. dimidiata*^39^. Infected nymphs of *R. prolixus* exhibited, on the contrary, a reduction of their locomotory activity^42^.

A low height device like the setup used in these experiments allowed us to highlight differences, between sexes, and between infected and non-infected insects. The questions that the results generate showed that little is known about the spatial distribution of the insects in their natural conditions and how they behave. Our results predict that in a natural/ anthropic environment the percentage of infected insects should increase with the height of the settlement. More experiments have to be carried out to understand the dispersion and aggregation behaviors of *T. infestans*, both in the laboratory and in the field. For example, one of our hypotheses concerns the influence of the height of the setup: the higher the setup, the greater would be the spatial segregation between the four categories. The response of the bugs should be modulated according to factors like the development stages and the physiological condition of the insects, the bug density, the numbers of available and suitable shelters, and the infection of the bugs. Another interesting question is about how mixed groups would distribute (both sexes, and/ or both infected and non-infected bugs). Indeed, in addition to those at work in monospecific aggregation new effects come into play among which segregation, whereby the different populations select different patches, plays a prominent role^59^.

Our results also open a new vision for controlling/ monitoring triatomines on the field, suggesting that there is a higher risk of *T. cruzi* infection in bugs located the upper part of walls/ rooms. More precise studies regarding bug distribution within microhabitats under field conditions should help to improve control and monitoring by trapping^60^. Finally, our results lead us to propose simple tests easily feasible in the field, based on geotaxis and the aggregative behavior of the bugs, to detect infected insects.

## Methods

*T. infestans* specimens were collected in dwellings from Yacuiba Municipality (Gran Chaco region), Department of Tarija, Bolivia, in the area Tierras Nuevas (S21.748334, W63.561866, 621 m asl) - San Francisco de Inti (S21.818193, W63.588042, 600 m asl). The infection rate of the captured insects, determined by analyzing drops of feces under a light microscope, was 47.5 ± 21.7%. Bugs were reared at 26±1°C, 60±15% RH, 12:12 night:dark cycle, in plastic pots containing a folded piece of kraft paper commonly used in the insectarium. They were fed on hens once every two weeks. Four conditions were then studied: non-infected males, non-infected females, infected males and infected females (abbreviated as NegMal, NegFem, PosMal, and PosFem in Figures). Fifteen and sixteen experiments were carried out with non-infected males and females respectively, and twelve and nine experiments with infected males and females respectively. Due to a problem in the insectarium, nine experiments using non-infected insects (4 and 5 experiments in males and females respectively) included some infected insects actually. A test for *T. cruzi* infection of the insects from these experiments was realized again determining a proportion of the infected individuals being less or equal to 20%. At 150 min, the aggregated fraction from the weakly infected experiments was closer to the fraction observed in non-infected group than to the fraction observed in infected groups (Anderson-Darling k-sample test: TkN = 6.08, P < 0.001; number of observations: 11 for non-infected males (NM) and for non-infected females (NF), 9 for infected females (IF), 12 for infected males (IM), 4 for weakly infected males (WIM) and 5 for weakly infected females (WIF); Anderson-Darling all-pairs comparison test: females: NF vs WIF: P = 0.45, WIF vs IF: P = 0.03; males: NM vs WIM: P = 0.91, WIM vs IM: P = 0.08). Therefore, these experiments were included in the non-infected group.

### Setup and Methods

A glass aquarium was used (50 × 20 × 50 cm) to avoid escaping of *T. infestans* which is unable to climb on glass walls. Insects were allowed to climb on one of the vertical surfaces of this aquarium (50 × 50 cm) offered by a paper sheet (kraft paper, 43 × 44 cm). The glass setup was washed, and the paper changed at the end of each experiment. It was illuminated by a centered 60W incandescent light bulb, placed at 50 cm behind the wall covered by the paper sheet. The paper guaranteed a homogeneous illumination of the setup. A video camera (Sony DCR-SR68) placed in front of the setup recorded the bug activity for 150 min. A 1 m high polystyrene wall surrounded the setup to isolate it. Experiments were conducted in a quiet and dark room to avoid any disturbance, at the beginning of the photophase. Ten bugs (8-10 days of starvation) were dropped on the bottom of the setup. They explored their environment rapidly and climbed on the wall. From the recordings, a snapshot was extracted at 1 min, 5 min and then every 5 min up to 150 min (31 snapshots in total). A processing program allowed us to record the spatial position of the thorax of each bug on each snapshot. With these spatial coordinates, the inter-individual distances were computed. As the length of an adult bug is on average 2.5 cm, and due to a tactile (legs or antennae) or visual perception, two individuals were considered as aggregated when they were at a distance less or equal to 4 cm.

### Indexes and statistics

Several indexes of position and aggregation were calculated using processing programs: 1) the number of individuals on the paper sheet; 2) the number of aggregated individuals; 3) the number and the size of the clusters; 4) the spatial stability of the individual (% of individuals that were found at time *t*+1 in a circle of 10mm in radius centered on the coordinate of the insect at time *t*); 5) the spatial stability of the biggest cluster (study of the distance between the centroid of the biggest cluster at time *t*+1 and the centroid of the biggest cluster at time *t*). Finally, the individual position of the insects in the setup at the end of each experiment was analyzed, recording the vertical orientation of the bugs (position 0: head towards the top), to put forward a privileged position. A vertical orientation was defined as inside an angle of ±30° to the vertical, head oriented towards the top or the bottom. Outside this range, the insect is not considered in a vertical position anymore. The structure of the clusters was also compared between infected males and infected females, conditions where bigger clusters emerged. Each cluster was considered as an undirected network where each node was an individual. Links between nodes were established when the distance between them was less or equal to 4 cm (the threshold for considering aggregation). The cluster K-density (ratio of the number of edges divided by the number of possible edges) was compared for clusters with a size greater than three individuals.

The comparisons between conditions were made using the Anderson-Darling k-sample test^61^. In case of obtaining a P < 0.05, an Anderson-Darling all-pairs comparison test was performed. These statistics were calculated using the functions adKSampleTest and adAllPairsTest of the *PMCMRplus* package of R^62,63^. Circular statistics were carried out with Oriana 4.02 (Kovach Computing services). Uniformity of data was tested using Rao’s test, mean comparisons using V-test, and distribution comparisons using Mardia-Watson-Wheeler pairwise test. The structure of the clusters was analyzed using the *igraph* package in R^64^. The linear regression was done using the lm function, and the comparison of the slopes of regression with the *lsmeans* package of R, using Least-squared means (predicted marginal means)^65^.

## Supporting information

## Acknowledgments

S. Depickère is particularly grateful to T. Chavez, F. Lardeux, Don Hugo and E. Siñani for logistic support. S. Depickère thanks the FYSSEN Foundation.

## Author contributions statement

SD and JLD designed the experiments. SD collected the data. SD, GMRA and JLD analyzed the results. SD and JLD wrote the main manuscript text. SD and GMRA prepared the figures. All authors reviewed the manuscript.

## Data availability statement

The datasets generated during and/or analysed during the current study are available from the corresponding author on reasonable request.

## References

1. WHO. Chagas disease (American trypanosomiasis). World Health Organization Fact Sheets 1 (2018).

2. Coura, J. R. & Albajar Viñas, P. Chagas disease: a new worldwide challenge. Nature 465, S6–7 (2010).

3. Rassi, A. J., Rassi, A. & Marin-Neto, J. A. Chagas disease. Lancet 375, 1388–1402 (2010).

4. Justi, S. A. & Galvão, C. The evolutionary origin of diversity in Chagas disease vectors. Trends Parasitol. 33, 42–52 (2017).

5. Flores-Ferrer, A., Marcou, O., Waleckx, E., Dumonteil, E. & Gourbière, S. Evolutionary ecology of Chagas disease; what do we know and what do we need? Evol. Appl. 11, 470–487 (2017).

6. Costa, J. T. The other insect societies. (Belknap Press of Harvard University Press, 2006).

7. Crall, J. D. et al. Social context modulates idiosyncrasy of behaviour in the gregarious cockroach *Blaberus discoidalis*. Anim. Behav. 111, 297–305 (2016).

8. Varadínová, Z., Stejskal, V. & Frynta, D. Patterns of aggregation behaviour in six species of cockroach: comparing two experimental approaches. Entomol. Exp. Appl. 136, 184–190 (2010).

9. Lazzari, C. R., Pereira, M. H. & Lorenzo, M. G. Behavioural biology of Chagas disease vectors. Mem. Inst. Oswaldo Cruz 108, 34–47 (2013).

10. Lardeux, F., Depickère, S., Duchon, S. & Chavez, T. Insecticide resistance of *Triatoma infestans* (Hemiptera, Reduviidae) vector of Chagas disease in Bolivia. Trop. Med. Int. Heal. 15, 1037–1048 (2010).

11. Depickère, S. et al. Susceptibility and resistance to deltamethrin of wild and domestic populations of *Triatoma infestans* (Reduviidae: Triatominae) in Bolivia: new discoveries. Mem. Inst. Oswaldo Cruz 107, 1042–1047 (2012).

12. Mougabure-Cueto, G. & Picollo, M. I. Insecticide resistance in vector Chagas disease: Evolution, mechanisms and management. Acta Trop. 149, 70–85 (2015).

13. Krause, J. & Ruxton, G. D. Living in groups. (Oxford University Press, 2002).

14. Depickère, S., Fresneau, D. & Deneubourg, J.-L. A basis for spatial and social patterns in ant species: dynamics and mechanisms of aggregation. J. Insect Behav. 17, 81–97 (2004).

15. Jeanson, R. et al. Self-organized aggregation in cockroaches. Anim. Behav. 69, 169–180 (2005).

16. Beard, C. Ben, Cordon-Rosales, C. & Durvasula, R. V. Bacterial symbionts of the Triatominae and their potential use in control of Chagas disease transmission. Annu. Rev. Entomol. 47, 123–141 (2002).

17. Schofield, C. J. The behaviour of Triatominae (Hemiptera: Reduviidae): a review. Bull. Entomol. Res. 69, 363–379 (1979).

18. Dambach, M. & Goehlen, B. Aggregation density and longevity correlate with humidity in first-instar nymphs of the cockroach (*Blattella germanica* L., Dictyoptera). J. Insect Physiol. 45, 423–429 (1999).

19. Yoder, J. A. & Grojean, N. C. Group influence on water conservation in the giant Madagascar hissing-cockroach, *Gromphadorhina portentosa* (Dictyoptera: Blaberidae). Physiol. Entomol. 22, 79–82 (1997).

20. Yoder, J. A., Hobbs, H. H. & Hazelton, M. C. Aggregate protection against dehydration in adult females of the cave cricket, *Hadenoecus cumberlandicus* (Orthoptera, Rhaphidophoridae). J. Cave Karst Stud. 64, 140–144 (2002).

21. Vitta, A. C. R., Lorenzo Figueiras, A. N., Lazzari, C. R., Diotaiuti, L. & Lorenzo, M. G. Aggregation mediated by faeces and footprints in *Triatoma pseudomaculata* (Heteroptera: Reduviidae), a Chagas disease vector. Mem. Inst. Oswaldo Cruz 97, 865–867 (2002).

22. Falvo, M. L., Lorenzo Figueiras, A. N. & Manrique, G. Spatio-temporal analysis of the role of faecal depositions in aggregation behaviour of the triatomine *Rhodnius prolixus*. Physiol. Entomol. 41, 24–30 (2015).

23. Reisenman, C. E., Lorenzo Figueiras, A. N., Giurfa, M. & Lazzari, C. R. Interaction of visual and olfactory cues in the aggregation behaviour of the haematophagous bug *Triatoma infestans*. J. Comp. Physiol. A 186, 961–968 (2000).

24. Rocha Pires, H. H. et al. Aggregation behaviour in *Panstrongylus megistus* and *Triatoma infestans:* inter and intraspecific responses. Acta Trop. 81, 47–52 (2002).

25. Cruz-Lopez, L., Malo, E. A. & Rojas, J. C. Aggregation pheromone in five species of Triatominae (Hemiptera: Reduviidae). Mem. Inst. Oswaldo Cruz 88, 535–539 (1993).

26. Minoli, S. A., Baraballe, S. & Lorenzo Figueiras, A. N. Daily rhythm of aggregation in the haematophagous bug *Triatoma infestans* (Heteroptera: Reduviidae). Mem. Inst. Oswaldo Cruz 102, 449–454 (2007).

27. Wigglesworth, V. B. The principles of insect physiology. (Chapman & Hall, 1972). doi: 10.1007/978-94-009-5973-6

28. Wisnivesky-Colli, C. et al. Laboratory comparison of feeding success among *Triatoma infestans, T. guasayana*, and *T. sordida* (Hemiptera□: Reduviidae). J. Med. Entomol. 32, 583–587 (1995).

29. Van Houte, S., Ros, V. I. & Van Oers, M. M. Walking with insects: molecular mechanisms behind parasitic manipulation of host behaviour. Mol. Ecol. 22, 3458–3475 (2013).

30. Hughes, D. P., Brodeur, J. & Thomas, F. (eds). Host Manipulation by Parasites. (Oxford University Press, 2012).

31. Hurd, H. Manipulation of medically important insect vectors by their parasites. Annu. Rev. Entomol. 48, 141–161 (2003).

32. Botto-Mahan, C. *Trypanosoma cruzi* induces life-history trait changes in the wild kissing bug *Mepraia spinolai*: implications for parasite transmission. Vector-Borne Zoonotic Dis. 9, 505–510 (2009).

33. Botto-Mahan, C., Cattan, P. E. & Medel, R. Chagas disease parasite induces behavioural changes in the kissing bug *Mepraia spinolai*. Acta Trop. 98, 219–223 (2006).

34. Lima, M. M., Borges-Pereira, J., Albuquerque Dos Santos, J. A., Teixeira Pinto, Z. & Vianna Braga, M. Development and reproduction of *Panstrongylus megistus* (Hemiptera: Reduviidae) infected with *Trypanosoma cruzi*, under laboratory conditions. Ann. Entomol. Soc. Am. 85, 458–461 (1992).

35. Takano-Lee, M. & Edman, J. D. Lack of manipulation of *Rhodnius prolixus* (Hemiptera: Reduviidae) vector competence by *Trypanosoma cruzi*. J. Med. Entomol. 39, 44–51 (2002).

36. Zeledón, R. El Triatoma dimidiata (Latreille, 1811) y su relación con la enfermedad de Chagas. (UNED, 1981).

37. Schaub, G. A. Developmental time and mortality of larvae of *Triatoma infestans* infected with *Trypanosoma cruzi*. Trans. R. Soc. Trop. Med. Hyg. 82, 94–97 (1988).

38. Oliveira, T. G. et al. Developmental and reproductive patterns of *Triatoma brasiliensis* infected with *Trypanosoma cruzi* under laboratory conditions. Mem. Inst. Oswaldo Cruz 105, 1057–1060 (2010).

39. Ramirez-Sierra, M. J., Herrera-Aguilar, M., Gourbière, S. & Dumonteil, E. Patterns of house infestation dynamic by non-domiciliated *Triatoma dimidiata* reveal a spatial gradient of infestation in rural villages and potential insect manipulation by *Trypanosoma cruzi*. Trop. Med. Int. Heal. 15, 77–86 (2010).

40. Nouvellet, P., Ramirez-Sierra, M. J., Dumonteil, E. & Gourbière, S. Effects of genetic factors and infection status on wing morphology of *Triatoma dimidiata* species complex in the Yucatán peninsula, Mexico. Infect. Genet. Evol. 11, 1243–1249 (2011).

41. Castro, L. A. et al. Flight behavior and performance of *Rhodnius pallescens* (Hemiptera: Reduviidae) on a tethered flight mill. J. Med. Entomol. 51, 1010–1018 (2014).

42. Marliére, N. P. et al. Trypanosomes modify the behavior of their insect hosts: effects on locomotion and on the expression of a related gene. PLoS Negl. Trop. Dis. 9, e0003973 (2015).

43. Lorenzo Figueiras, A. N. & Lazzari, C. Aggregation behavior and interspecific responses in three species of Triatominae. Mem. Inst. Oswaldo Cruz 93, 133–137 (1998).

44. Lorenzo Figueiras, A. N., Kenigsten, A. & Lazzari, C. R. Aggregation in the haematophagous bug *Triatoma infestans:* chemical signals and temporal pattern. J. Insect Physiol. 40, 311–316 (1994).

45. Tanaka, S., Wolda, H. & Denlinger, D. L. Group size affects the metabolic rate of a tropical beetle. Physiol. Entomol. 13, 239–241 (1988).

46. Benoit, J. B. et al. Mechanisms to reduce dehydration stress in larvae of the Antarctic midge, *Belgica antarctica*. J. Insect Physiol. 53, 656–667 (2007).

47. Schaub, G. A. Does *Trypanosoma cruzi* stress its vectors? Parasitol. Today 5, 185–188 (1989).

48. Silverman, J. M. & Bell, W. J. The role of vertical and horizontal object orientation in mate-finding and predator-avoidance by the American cockroach. Anim. Behav. 27, 652–657 (1979).

49. Russomando, G. et al. *Trypanosoma cruzi:* polymerase chain reaction-based detection in dried feces of *Triatoma infestans*. Exp. Parasitol. 83, 62–66 (1996).

50. Marcet, P. L. et al. PCR-based screening and lineage identification of *Trypanosoma cruzi* directly from faecal samples of triatomine bugs from northwestern Argentina. Parasitology 132, 57–65 (2006).

51. Peterson, J. K., Graham, A. L., Dobson, A. P. & Chavez, O. T. *Rhodnius prolixus* life history outcomes differ when infected with different *Trypanosoma cruzi* I strains. Am. J. Trop. Med. Hyg. 93, 564–572 (2015).

52. Hughes, D. P., Kathirithamby, J., Turillazzi, S. & Beani, L. Social wasps desert the colony and aggregate outside if parasitized□: parasite manipulation? Behav. Ecol. 15, 1037–1043 (2004).

53. Lefèvre, T. & Thomas, F. Behind the scene, something else is pulling the strings: Emphasizing parasitic manipulation in vector-borne diseases. Infect. Genet. Evol. 8, 504–519 (2008).

54. Jacquin, L., Mori, Q., Steffen, M. & Medoc, V. Non-specific manipulation of gammarid behaviour by *P. minutus* parasite enhances their predation by definitive bird hosts. PLoS One 9, e101684 (2014).

55. Arnal, A. et al. Activity level and aggregation behavior in the crustacean gammarid *Gammarus insensibilis* parasitized by the manipulative trematode *Microphallus papillorobustus*. Front. Ecol. Evol. 3, 109 (2015).

56. Weinersmith, K. L. et al. *Euhaplorchis californiensis* cercariae exhibit positive phototaxis and negative geotaxis. J. Parasitol. 104, 329–333 (2018).

57. Thomas, F., Adamo, S. & Moore, J. Parasitic manipulation: where are we and where should we go? Behav. Processes 68, 185–199 (2005).

58. Lazzari, C. R. & Lorenzo, M. G. Exploiting triatomine behaviour: alternative perspectives for their control. Mem. Inst. Oswaldo Cruz 104, 65–70 (2009).

59. Nicolis, S. C., Halloy, J. & Deneubourg, J.-L. Transition between segregation and aggregation: the role of environmental constraints. Sci. Rep. 6, 32703 (2016).

60. Rojas de Arias, A. et al. Post-control surveillance of *Triatoma infestans* and *Triatoma sordida* with chemically-baited sticky traps. PLoS Negl. Trop. Dis. 6, e1822 (2012).

61. Scholz, F. & Stephens, M. K-Sample Anderson-Darling Tests. J. Am. Stat. Assoc. 82, 918–924 (1987).

62. R Core Team. R: A language and environment for statistical computing. (R Foundation for Statistical Computing, 2016).

63. Pohlert, T. PMCMRplus: calculate pairwise multiple comparisons of mean rank sums extended. R Packag. version 1.0.1 (2018).

64. Csardi, G. & Nepusz, T. The igraph software package for complex network research. Inter Journal **Complex Sy**, 1695 (2006).

65. Lenth, R. V. Least-Squares Means: the R package lsmeans. J. Stat. Softw. 69, 1:33 (2016).

