## Supplementary material for "Alteration of the social and spatial organization of the vector of Chagas disease, *Triatoma infestans*, by the parasite *Trypanosoma cruzi*"

**Figure S1. Examples of insect's positions at 150 min in the setup for the four conditions**

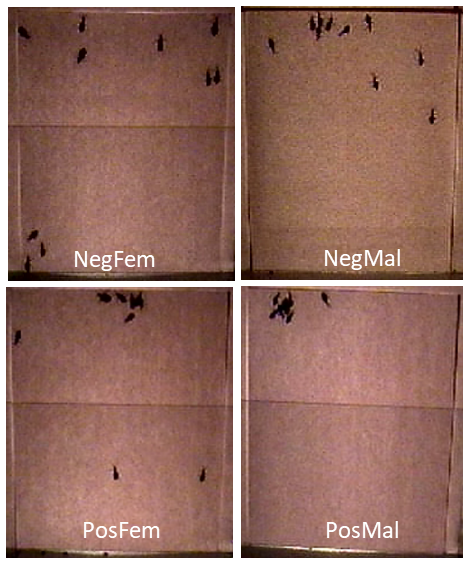

**Figure S2. Boxplot distribution of the median fraction of individuals at the top of the setup (40-44cm) at 150 min**. Number of observations: 16, 15, 9, 12 for NegFem, NegMal, PosFem, and PosMal respectively. Anderson-Darling k-sample test between the four conditions for all individuals: TkN = 8.0, P < 0.001. Results of Anderson-Darling all-pairs comparison tests are shown at the top of the figure (conditions with different letters correspond to conditions statistically different at P < 0.04)

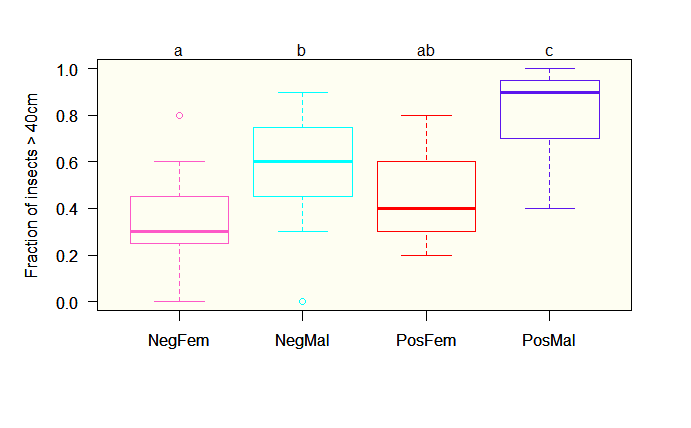

**Figure S3. Relative frequency of experiments according to the aggregated fraction of individuals for the four conditions**

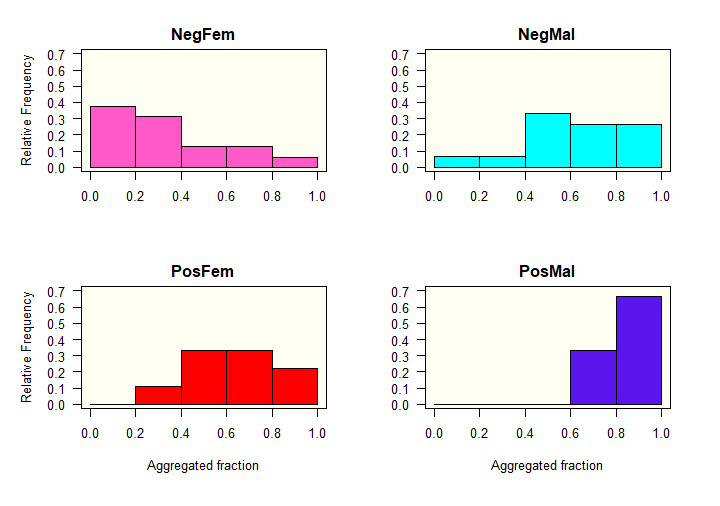

**Figure S4. Example of clusters of 10 bugs in infected conditions**. Nodes represent the insect positions on the wall. Links between nodes are present only when the distance is less or equal to 4 cm (the aggregative distance)

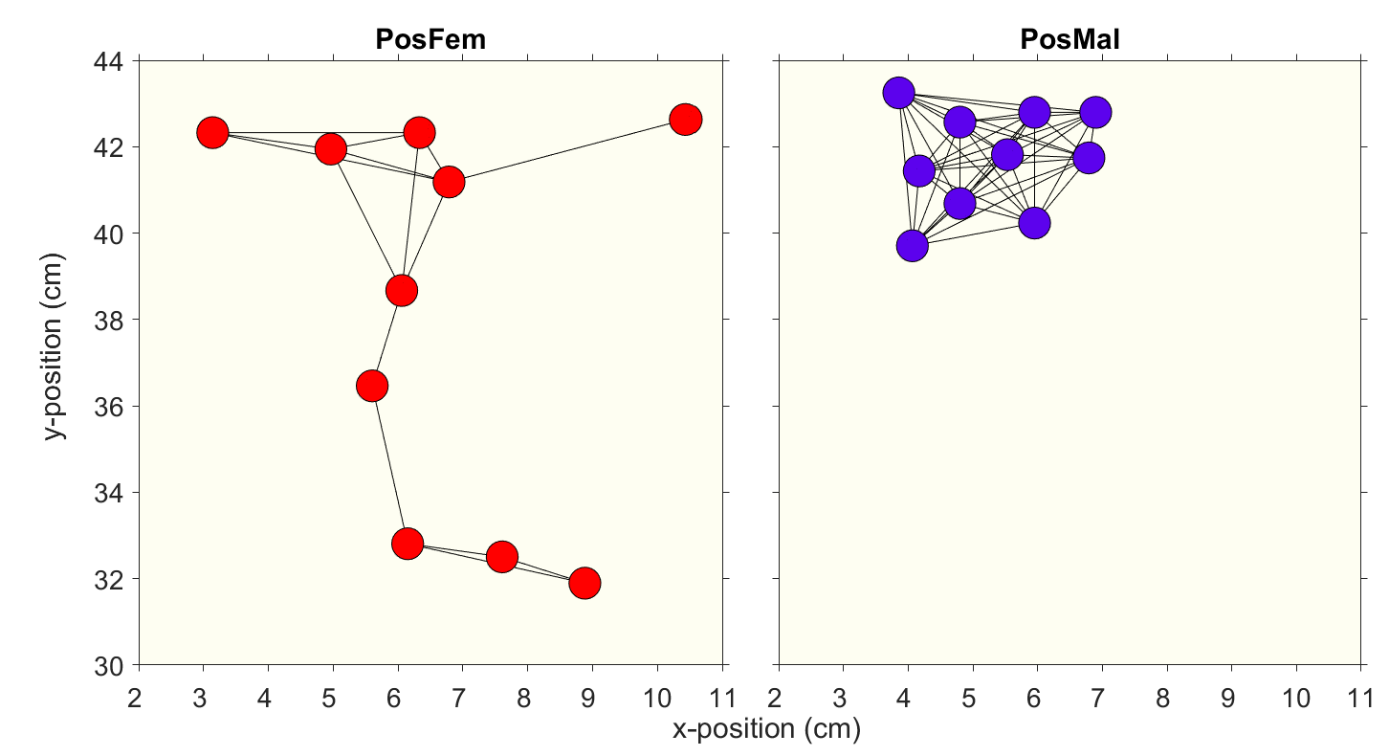

**Figure S5. Vertical distribution of insects on the wall for the four conditions**. Fraction of bugs by horizontal strips of 5 mm, from 0 to 440 mm, corresponding to the height of the setup for non-infected (females: pink, males: cyan) and infected bugs (females: red, males: blue)

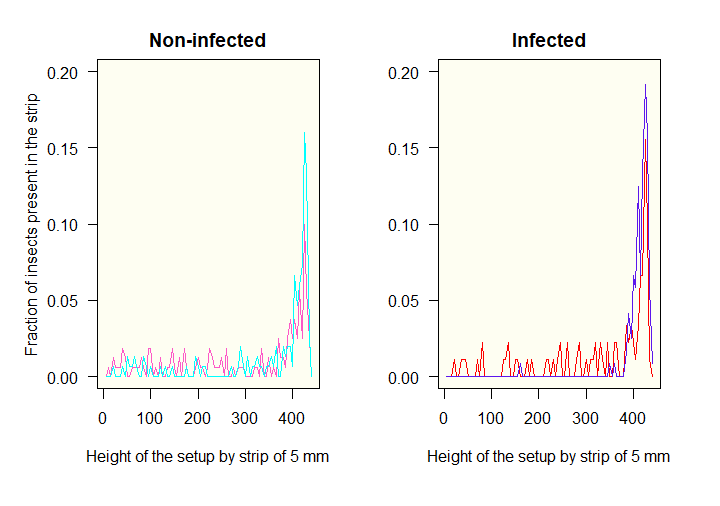
